# EpiCHAOS: a metric to quantify epigenomic heterogeneity in single-cell data

**DOI:** 10.1101/2024.04.24.590899

**Authors:** Katherine Kelly, Michael Scherer, Martina Maria Braun, Pavlo Lutsik, Christoph Plass

## Abstract

Epigenetic heterogeneity is a fundamental property of biological systems, and is recognized as a potential driver of tumor plasticity and therapy resistance. Single-cell epigenomics technologies have been widely employed to study epigenetic variation between – but not within – cellular clusters. We introduce epiCHAOS: a quantitative metric of cell-to-cell heterogeneity, applicable to any single-cell epigenomics data type. After validation in synthetic datasets, we applied epiCHAOS to investigate global and region-specific patterns of epigenetic heterogeneity across diverse biological systems. EpiCHAOS provides an excellent approximation of stemness and plasticity in development and malignancy, making it a valuable addition to single-cell cancer epigenomics analyses.

## Background

Cell-to-cell heterogeneity can be found at multiple levels in all complex biological systems – ranging from that within populations of genetically diverse individuals, to the non-genetic functional heterogeneity between tissues and cell types, and more subtle molecular differences between cells within the same cell type^1^. In recent years this phenomenon of inter-cellular heterogeneity has received particular attention in the study of cancer as a potential driver of tumor progression and therapy resistance ^2,3,4^.

Although cancer formation begins with the clonal expansion of a single transformed cell, most tumors ultimately acquire extensive heterogeneity, with a single tumor consisting of several populations of cells with diverse phenotypic characteristics ^2,5^. This is partly due to the emergence of genetically distinct subclones. However, epigenetic dysregulation is now also recognized as a hallmark of cancer, and it is becoming increasingly evident that profound functional heterogeneity can arise within genetically identical cellular structures due to changes in the epigenetic landscape, including DNA and histone modifications and the larger chromatin architecture ^6,7^.

Outside of malignancy, recent studies have highlighted the essential role of cellular heterogeneity in maintaining pluripotency and in shaping differentiation trajectories in the developing organism ^8^. Generally, multipotency is accompanied by increased stochastic molecular variation or “noise” which is lost as cells differentiate and commit to a specific cell fate, in which a more stable transcriptional program is acquired ^9^. In the context of malignancy, such regulatory heterogeneity or “noise” could similarly act as a driving force towards functional intratumor heterogeneity ^9^. Here epigenetic heterogeneity could have potentially detrimental implications. By creating a fitness advantage and improving a tumor’s ability to adapt to a variety of intrinsic and extrinsic stresses, heterogeneity increases the chances that part of the tumor will tolerate a range of therapeutic insults, and thus represents a major clinical challenge ^5^. Moreover, such heterogeneity can allow the “division of labor” required for a tumor to function efficiently as a system. For example, the functional diversity required to undertake the metastatic cascade – involving degradation of extracellular matrix, invasion into local tissues, survival in transit through the bloodstream and ultimate colonization in a distant organ – is an unthinkable feat for a homogeneous population of cells, but is likely accomplished by a system of phenotypically heterogeneous cells, with each taking on different tasks to the benefit of the whole ^10^ ^11^.

An appreciation of intratumor heterogeneity is therefore essential to the study of cancer, however this has historically been overlooked due to the inability of traditional bulk sequencing approaches to unmask intrinsic variations within cell populations. Since the advent of single-cell sequencing technologies, it is now possible to disentangle intratumor and microenvironmental heterogeneity at unprecedented resolution at the genomic, transcriptomic and epigenomic levels ^12^. Using technologies such as the Assay for Transposase Accessible Chromatin with sequencing (scATAC-seq), the presence of epigenetically distinct clusters within tumors – the simplest layer of epigenetic heterogeneity – has been demonstrated across a broad range of cancer types ^13,14,15,16^. However, a more complex and underexplored problem is quantifying heterogeneity between cells within a given group or cluster.

A few recent studies have incorporated methods to quantify cell-to-cell heterogeneity at the transcriptional level using distance-based or entropy-based metrics, or to quantify transcriptional “noise” using metrics such as the coefficient of variation ^17–19,20^. Others have taken advantage of read-level DNA methylation data to devise metrics of epigenetic heterogeneity that can be applied to bulk datasets ^21–25^. However, such strategies have not been extended to single-cell applications and are therefore limited in discerning heterogeneity *between* from heterogeneity *within* cellular clusters. A metric for quantifying epigenetic heterogeneity in single-cell datasets has so far not been established.

To address this we developed epiCHAOS (<u>Epi</u>genetic/<u>C</u>hromatin <u>H</u>eterogeneity <u>A</u>ssessment <u>O</u>f <u>S</u>ingle cells), a distance-based heterogeneity score designed to quantify cell-to-cell epigenetic heterogeneity using single-cell epigenomic data. After systematically validating epiCHAOS in a range of synthetic and real datasets, we demonstrated its ability to capture features of stemness and plasticity in development and malignancy. Additionally, we employed epiCHAOS for investigating region-specific and pathway-specific differences in epigenetic heterogeneity, and demonstrated its applicability to a range of single-cell epigenomic data types.

## Results

### EpiCHAOS reliably quantifies epigenetic heterogeneity in single-cell epigenomics data

We designed epiCHAOS to assign a heterogeneity score at the level of cell clusters, or other user-defined groups of interest e.g. cell types or treatment conditions. Here, we initially focused on scATAC-seq data as the most commonly used single-cell epigenomics modality. To establish epiCHAOS scores, data from each cluster were extracted as a binarized matrix, representing a peaks-by-cells or tiles-by-cells matrix in the case of scATAC-seq. For each cluster we then computed the distances between all pairs of cells using a count-centered version of the Jaccard distance ^26^, and then took the mean of all pairwise distances per cluster as its epiCHAOS score (Figure 1A).

**Figure 1.**
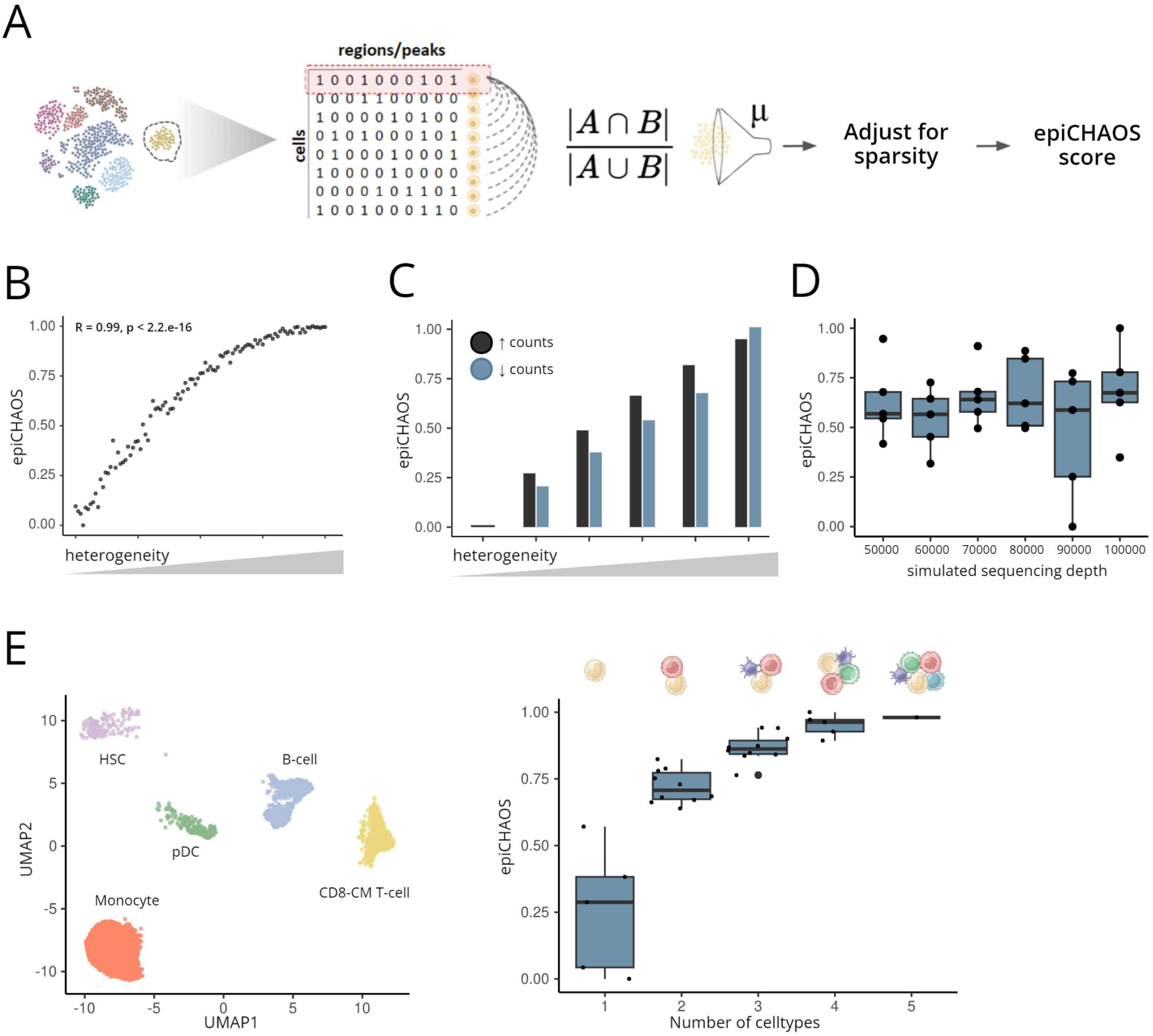
EpiCHAOS reliably quantifies epigenetic heterogeneity in single-cell epigenomics data. **A.** Schematic describing epiCHAOS calculation. Using single-cell epigenomics data in binarized matrix form, epiCHAOS scores are assigned per cluster by computing the mean of all pairwise cell-to-cell distances using a chance-centered Jaccard index followed by regression-based adjustment for sparsity. μ=mean per cluster. **B.** Scatter plot illustrating the correlation between epiCHAOS scores (epiCHAOS) and controlled heterogeneity across 100 synthetic datasets. Pearson correlation coefficient and p-value is shown. **C.** Barplots illustrate increasing heterogeneity after perturbation of scATAC-seq data from sorted monocytes by either randomly adding or randomly removing 10-50% of 1’s. **D.** Boxplot comparing epiCHAOS scores across six simulated single-cell ATAC-seq datasets with varying sequencing depths. Data were simulated using scReadSim with sequencing depth varying from 50,000 to 100,000 counts. ScATAC-seq data from hematopoietic stem cells subset from the Granja *et al.* 2019 dataset ^28^ were used as the baseline counts matrix. **E.** Validation of epiCHAOS using *in silico* mixtures of hematopoietic cell types. UMAP embedding illustrates scATAC profiles from five selected cell types of human bone marrow ^28^. After selecting 500 top differentially accessible peaks for each cell type, *in silico* mixtures of two to five cell types in all possible combinations were created. Boxplots show the relationship between epiCHAOS scores (epiCHAOS) and number of cell types (y-axis) after *in silico* mixing.

To validate the design of epiCHAOS we generated a range of fully synthetic scATAC-seq datasets in which we controlled the levels of heterogeneity, while maintaining a stable total count (see “Methods” for details). We showed that the epiCHAOS score is highly correlated with the true controlled, artificial heterogeneity (Pearson R=0.99) (Figure 1B). Next, we verified that heterogeneity can be detected both in cases of increasing and decreasing counts representing a genome-wide loss/gain of chromatin accessibility. To achieve this, we simulated a series of scATAC-seq datasets in which we incrementally increased heterogeneity, while either increasing or decreasing genome-wide chromatin accessibility (see “Methods” for details). We showed that epiCHAOS correctly detects differences in heterogeneity both in cases where counts are added and removed (Figure 1C).

To confirm that epiCHAOS does not perceive differences in sparsity as differences in heterogeneity, we generated a series of random datasets with varying total numbers of 1’s. We found no correlation between epiCHAOS scores and total number of 1’s between datasets (Figure S2A). In real datasets however, differences in coverage may be more complex since they are accompanied by differences in the number of missing values representing false negatives. Thus, we found that higher heterogeneity can be perceived in data with lower total number of fragments (Figure S2B). Genome-wide differences in detected ATAC signals can arise from both technical or biological reasons, for example due to the presence of quiescent cells or in cells with copy number alterations (CNAs) (Additional file 1: Figure S1A-D), or due to differences in sequencing depth. To exclude any effect of this on epiCHAOS scores we implemented a linear regression-based adjustment for the genome-wide chromatin accessibility across cell clusters, which gave us a count-adjusted heterogeneity score. We showed that this adjusted score is no longer affected by differences in genome-wide chromatin accessibility and is robust to the presence of large-scale deletions and gains in tumor samples (Additional file 1: Figure S1A-D). To assess the effect of sequencing depth, we used the published tool, scReadSim^27^. We found that epiCHAOS was not affected by differences in sequencing depth (Figure 1D).

We further validated epiCHAOS using *in silico* mixtures of cell types from the human hematopoietic system. We selected five distinct cell types from a previously published hematopoietic dataset ^28^ including hematopoietic stem cells (HSCs), monocytes, B-cells, CD8-T cells and plasmacytoid dendritic cells (pDCs). After reducing the peaks matrix to regions which are most differentially accessible between cell types (500 peaks for each cell type), we created mixtures of two to five cell types in all possible combinations and applied epiCHAOS to each individual cell type and mixture. As expected, epiCHAOS scores were relatively low in individual cell types and increased with the number of cell types in the mixture (Figure 1E). We also confirmed that epiCHAOS was not influenced by technical noise or other potential technical confounders (Additional file 1: Figure S1E-H, Additional file 2: Supplementary Note 1), and is minimally influenced by differences in clustering resolution (Additional file 1: Figure S3, Figure S4, Additional file 2: Supplementary Note 2).

### EpiCHAOS reflects epigenetic heterogeneity associated with developmental plasticity

In order to show that epiCHAOS is a reliable metric that leads to biologically plausible results, we initially focused on applications where there was some prior knowledge or expectation. For example, previous studies have recognized epigenetic heterogeneity as a feature of uncommitted, multipotent cell states, which decreases in more committed, differentiated cells ^9, 29^. To test if epiCHAOS scores align with this expectation we applied our score to scATAC-seq data from a range of developmental contexts. First, we tested epiCHAOS in the human hematopoietic system^28,29^, where we detected highest epigenetic heterogeneity in progenitor cells including hematopoietic stem cells (HSCs), Common Lymphoid Progenitors (CLPs), Common Myeloid Progenitors (CMPs), Lympho-myeloid primed progenitor cells (LMPPs), and early erythroid cells. In contrast, more differentiated cells of the myeloid, erythroid and lymphoid lineages showed lower epiCHAOS scores (Figure 2A). Second, in data from mouse gastrulation^30^, epiCHAOS scores were highest in less differentiated cells. This was especially notable in the primitive streak – the point at which cells of the epiblast undergo epithelial-to-mesenchymal transition (EMT) – and in primordial germ cells. Low epiCHAOS scores were detected throughout the formation of distinct meso-, endo- and ectodermal lineages (Figure 2B). In Drosophila embryogenesis, we also observed high epigenetic heterogeneity at the earliest multipotent stages including undifferentiated cells, blastoderm and germ cells (Figure 2C). Among more differentiated tissues, we detected high heterogeneity scores in the neuronal compartments, likely due to their extraordinary functional heterogeneity^31^. We confirmed using the Descartes scATAC-seq tissue atlas that higher heterogeneity in neural tissues persists outside of an embryonic context ^32^ (Additional file 1: Figure S5).

**Figure 2.**
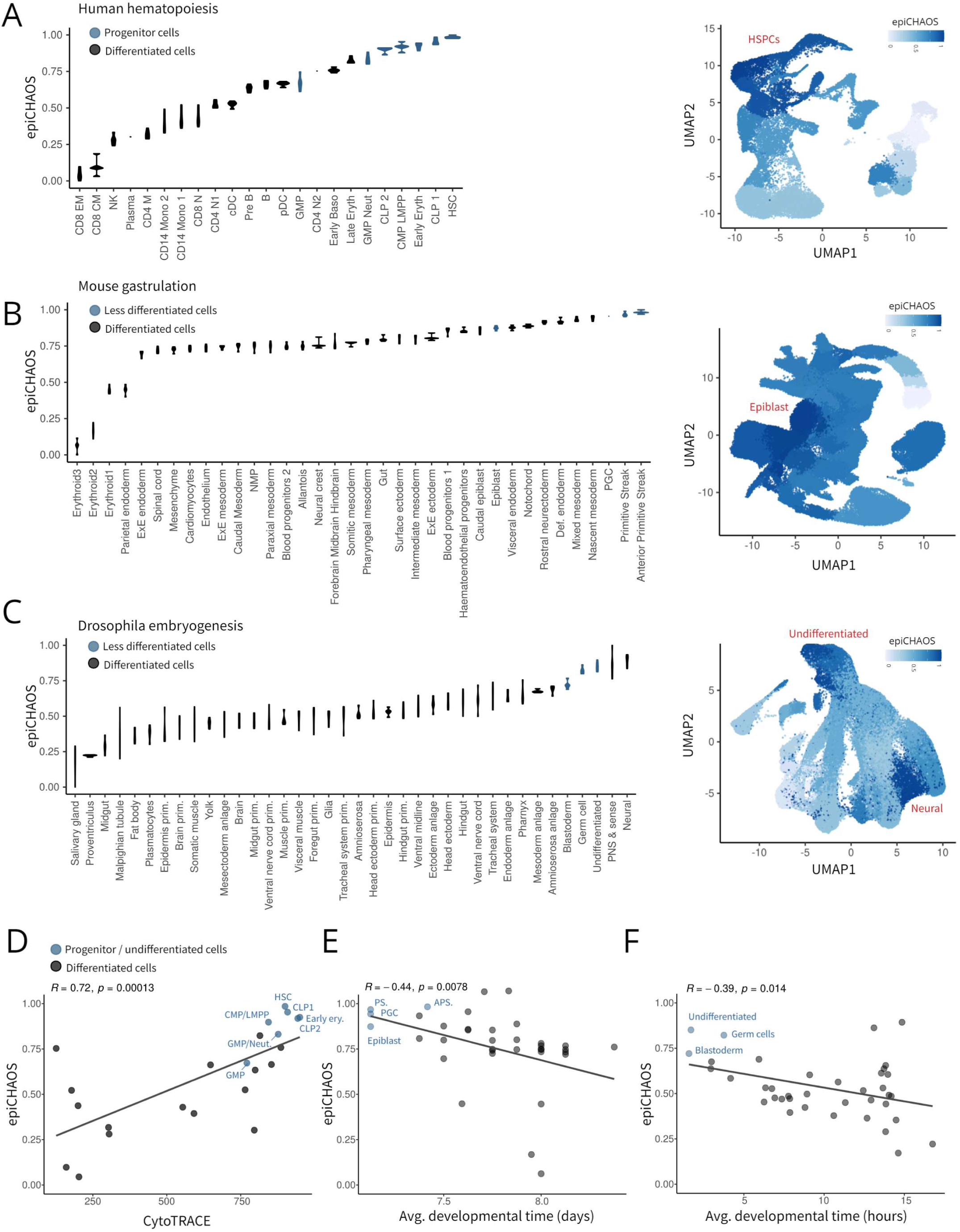
EpiCHAOS reflects epigenetic heterogeneity associated with developmental plasticity. **A-C.** Violin plots (left) showing epiCHAOS scores (epiCHAOS) computed in scATAC-seq data from **A.** human hematopoiesis^28^, **B.** mouse gastrulation^30^ and **C.** drosophila embryogenesis^55^. EpiCHAOS scores were computed per-cell type as annotated in the original publications. Violins represent the scores computed in five random subsamples of 100 cells from each cell type, or once where fewer than 100 cells were available. Plots are ordered by epiCHAOS scores and progenitor cells and undifferentiated tissue types are highlighted in blue. UMAP embeddings (right) illustrating epiCHAOS scores in the same datasets as in violin plots. UMAPs are colored by epiCHAOS scores computed per annotated cell/tissue type. **D-F.** Scatter plots correlating epiCHAOS scores (epiCHAOS) with developmental time as defined by (D) CytoTRACE score, averaged across cells (human hematopoietic system), (E) developmental time in days at sample collection, averaged across cells (mouse gastrulation), or (F) predicted developmental time in hours (Drosophila embryogenesis). EpiCHAOS scores represent the average of pseudo-replicates shown in A-C. Linear regression lines are displayed with Pearson correlation coefficients and p-values.

To validate these observations we correlated our score with CytoTRACE^33^ – a scRNA-based metric designed to capture stemness/plasticity – using scRNAseq data from hematopoietic cells^28^. EpiCHAOS correlated moderately with CytoTRACE scores (Figure 2D), in some cases following a pattern better representative of the differentiation trajectory than CytoTRACE. For example, epiCHAOS detected higher heterogeneity in naive CD8+ and CD4+ T-cells compared to memory T cells. Naive T cells are expected to be more developmentally plastic, but are not detected as such by CytoTRACE. This suggests that epiCHAOS captures some features of cellular plasticity that might not be detectable at the transcriptional level. Similarly, in gastrulation and embryogenesis datasets, epiCHAOS correlated with previously annotated metrics of developmental time (Figure 2E-F). This pattern of epigenetic heterogeneity was also in most cases reflected by higher transcriptional heterogeneity in less differentiated cells (Additional file 1: Figure S6).

Collectively these data suggested that epiCHAOS can provide an accurate approximation of epigenetic heterogeneity indicative of developmental plasticity.

### EpiCHAOS correlates with features of malignant cell plasticity and aging

To investigate epigenetic heterogeneity in malignancy we selected two previously published scATAC-seq datasets from 16 breast^13^ and 16 liver^34^ cancer patients. We subsetted each dataset for epithelial cells and applied epiCHAOS to the resulting epithelial clusters (Figure 3A, Additional file 1: Figure S7A-B). To understand what features of tumor cells coincide with higher epigenetic heterogeneity, we assigned molecular signature scores to each cluster using the scATAC-seq gene score matrices. Within each tumor type we correlated epiCHAOS against each molecular signature.

**Figure 3.**
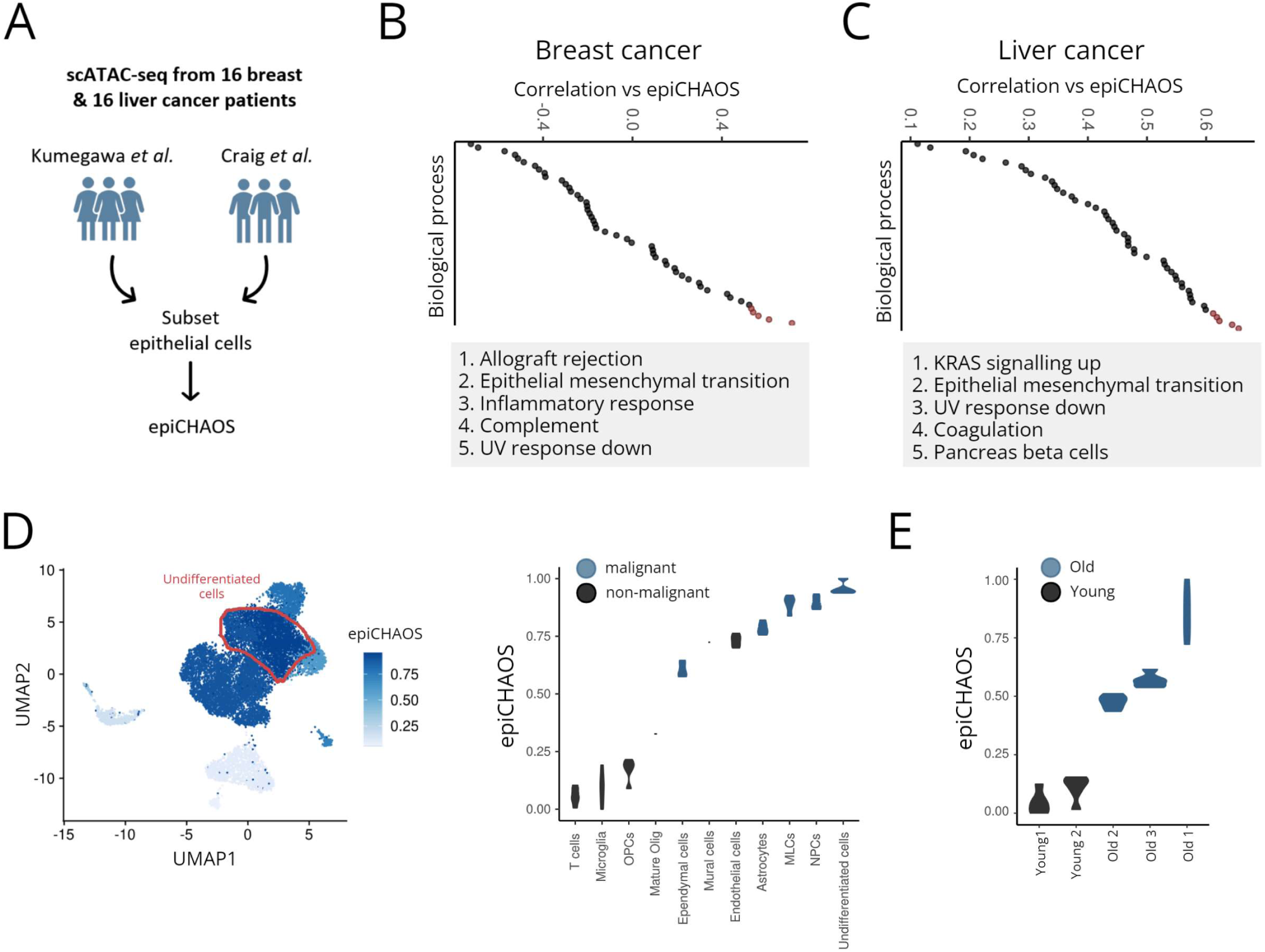
EpiCHAOS correlates with features of malignant cell plasticity and aging. **A.** Schematic describing breast and liver cancer datasets used for epiCHAOS calculation. **B-C.** Dot plots illustrate ordered Pearson coefficients after correlation of per-cluster epiCHAOS scores against gene set scores for all Hallmark Gene Ontology biological processes in breast (B) and liver cancer (C). Top 5 correlations in each dataset are highlighted and labeled below. **D.** UMAP embedding (left) of scATAC-seq data from five primary and two metastatic childhood ependymoma tumors^36^. Cells are colored by epiCHAOS scores (epiCHAOS) computed for each cell type. Undifferentiated cells are highlighted in red. Violin plot (right) ordered by epiCHAOS scores (epiCHAOS) for all malignant and non-malignant cell types annotated in ependymoma tumors. Malignant cell types are highlighted in blue. Violins represent the scores computed in five random subsamples of 100 cells from each cell type, or once where fewer than 100 cells were available. **E.** Violin plot ordered by epiCHAOS scores computed in scATAC-seq data from old (blue; n = 3) and young (black; n = 2) mouse HSCs^39^. Violins represent the scores computed in five random subsamples of 100 cells from each group.

In breast cancer, we noticed that epiCHAOS scores were highly correlated with gene sets relating to EMT – a process which is considered integral to breast cancer plasticity and metastasis^35^ (Figure 3B). EMT-related signaling pathways such as TGF-beta and WNT signaling were also correlated with epiCHAOS scores (Additional file 1: Figure S7C). By contrast, gene sets related to estrogen receptor signaling, which are associated with better breast cancer prognosis, were anticorrelated with epiCHAOS scores (Additional file 3: Table S7). Clusters with high epiCHAOS scores also had higher accessibility at genes within various previously described gene expression signatures of the more aggressive invasive and metaplastic breast cancer subtypes (Additional file 1: Figure S7C). Similarly, in liver cancer, EMT-related gene sets as well as previously derived liver cancer prognostic signatures were increased in clusters with high epiCHAOS scores (Figure 3C, Additional file 1: Figure S7C).

To support a suspected link between epigenetic heterogeneity and plasticity/stemness we applied epiCHAOS to scATAC-seq data from childhood ependymoma comprising multiple differentiated (astrocytes, ependymal cells, neural progenitor cells, and mesenchymal-like cells) as well as undifferentiated tumor cell types^36^. Here, we detected the highest epiCHAOS scores in undifferentiated cell populations, which are known to be enriched in more aggressive ependymomas^37,38^ (Figure 3D). Among malignant cell types, the lowest heterogeneity scores were detected in ependymal cells (Figure 3D), which represent the most differentiated ependymoma tumor cells and which are associated with less aggressive disease^37^.

We also wondered whether epiCHAOS would detect increased epigenetic heterogeneity associated with organism aging. Using a scATAC-seq dataset from young (2 months) and old (24 months) mouse HSCs^39^, epiCHAOS detected a global increase in epigenetic heterogeneity in the aged cells (Figure 3E, Additional file 3: Table S10), reflecting the stochastic loss of epigenetic information which is considered a hallmark of aging^40^.

Overall these data support a connection between epigenetic heterogeneity quantified with epiCHAOS and cancer cell plasticity and aggressiveness, as well as aging in the hematopoietic system.

### EpiCHAOS reveals elevated levels of epigenetic heterogeneity at PRC2 targeted regions and promoters of developmental genes

Beyond comparisons of global epigenetic heterogeneity between cell states, we also wanted to apply epiCHAOS to investigate differences in epigenetic heterogeneity at various classes of genomic loci.

Focusing on the hematopoietic system, we utilized annotated chromatin/transcription factor binding sites from the Encyclopedia of DNA Elements (ENCODE) database to investigate differences in heterogeneity across different genomic regions. We found that epiCHAOS scores were especially high at Polycomb Repressive Complex 2 (PRC2) targeted regions, as well as at binding sites for CCCTC-binding factor (CTCF) and Cohesin (Figure 4A, Additional file 3: Table S11). Per-region heterogeneity scores tended to be highly correlated between different hematopoietic cell types (Additional file 1: Figure S8A). We validated this finding by comparison to DNA methylation variation in hematopoietic cells as an independent measure of epigenetic heterogeneity. At the DNA methylation level, epigenetic heterogeneity was similarly highest at PRC2 targets, while binding sites for CTCF and Cohesin were also among the most variable regions (Figure 4B). Visualization of peaks-by-cells matrices at examples of regions with high and low epiCHAOS scores reflected the reported differences in heterogeneity (Figure 4C). To understand whether this finding translates to the transcriptional level, we also calculated transcriptional noise for each gene in HSCs based on the coefficient of variation. Consistent with previous reports^41,8^, and in line with their epigenetic heterogeneity, we found that transcriptional noise was significantly higher at PRC2 targeted genes compared to non-PRC2 targets (Additional file 1: Figure S9A).

**Figure 4.**
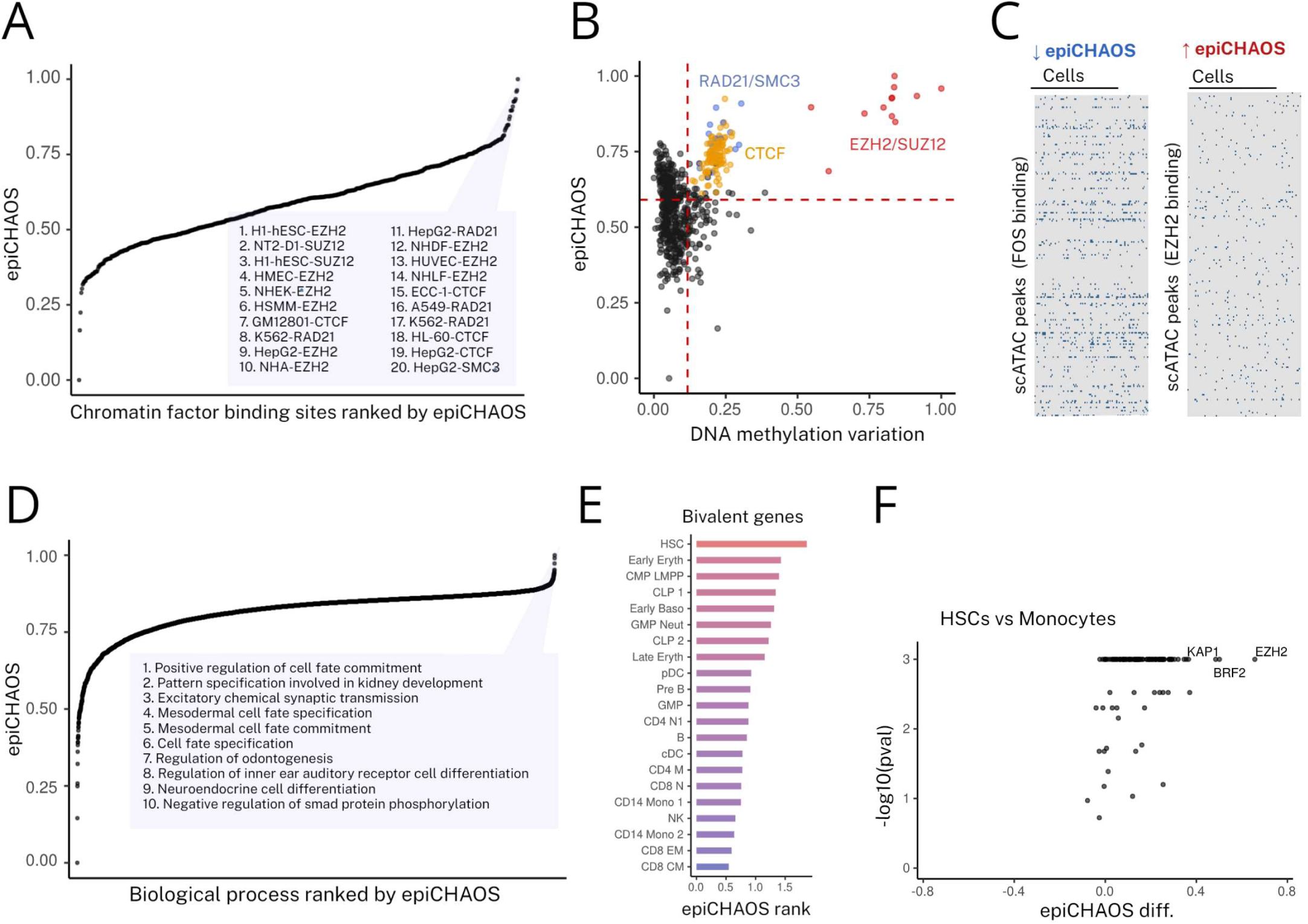
EpiCHAOS reveals elevated levels of epigenetic heterogeneity at PRC2 targeted regions and promoters of developmental genes. **A.** Ordered dot plot showing epiCHAOS scores (epiCHAOS) computed across region sets for each ENCODE chromatin factor binding site in hematopoietic stem cells, ordered by epiCHAOS scores. Top 20 region sets (cell type and binding site) are labeled. **B.** Scatter plot comparing epiCHAOS scores (epiCHAOS) with DNA methylation variation at each transcription factor binding site as in (A). PRC2 targets (binding sites for EZH2/SUZ12, red), CTCF targets (orange) and Cohesin binding sites (binding sites for RAD21/SMC3, blue) are highlighted. X-axis represents the average of per-CpG methylation variances across 10 individuals at all CpGs overlapping with the respective region set. Values are scaled to a 0-1 range. **C.** Heatmaps show examples of high-epiCHAOS (EZH2 binding sites) and low-epiCHAOS (FOS binding sites) regions in the peaks matrix of scATAC-seq data from HSCs. **D.** Ordered dot plot showing epiCHAOS scores (epiCHAOS) computed across promoters for each gene ontology biological process in hematopoietic stem cells, ordered by epiCHAOS scores. **E.** Bar plot comparing epiCHAOS ranks for the set of bivalent genes across different hematopoietic cell types. The higher the rank indicates that the selected gene set has higher epiCHAOS scores compared to other gene sets in that celltype. Ranks were -log(10) transformed for display. **F.** Volcano plot illustrates differential heterogeneity between hematopoietic stem cells and monocytes. Differential heterogeneity was tested for each ENCODE TFBS (binding sites measured in K562 cells). For each TFBS, the -log10(p-value) obtained by permutation test is displayed on the y-axis, and the difference in epiCHAOS scores between the two cell types is displayed on the x-axis.

To further understand the characteristics of regions displaying high epigenetic heterogeneity, we computed epiCHAOS scores at promoter-associated scATAC-seq peaks over a range of gene sets. Gene sets with the highest heterogeneity scores were enriched for developmental processes such as cell fate commitment, cell fate specification and somatic stem cell division (Figure 4D, Additional file 3: Table S12). Per-gene-set heterogeneity scores were correlated between cell types (Additional file 1: Figure S8B), however certain gene sets showed more cell-type specific heterogeneity patterns. For example, we noted that genes related to cell fate specification which were among the most heterogeneous gene sets in HSCs, ranked lower in heterogeneity in more differentiated cell types (Additional file 1: Figure S9B). Similarly, promoters of bivalent genes ranked among the most heterogeneous gene sets in HSCs and less so in more differentiated cell types (Figure 4E).

We wondered whether certain genomic regions contributed to the differences in heterogeneity we had observed between hematopoietic stem/progenitors and more differentiated cell types. Comparing HSCs to monocytes we found that the largest increase in heterogeneity in HSCs was at Enhancer Of Zeste 2 (EZH2) binding sites, followed by BRF2 and KAP1 binding sites (Figure 4F). Comparison of HSCs to differentiated B-cells and CD8 T-cells yielded a similar pattern, suggesting that epigenetic heterogeneity at these key genomic regions might contribute to maintaining pluripotency (Additional file 1: Figure S9C-D).

### EpiCHAOS is applicable to single-cell epigenomics data from any modality

After demonstrating epiCHAOS’ capability in scATAC-seq data, we wanted to test its application to other epigenomics layers. For this we first applied epiCHAOS to single-cell nucleosome, methylation and transcription sequencing (scNMT-seq) DNA methylation data from mouse gastrulation^42^. Here we observed highest epigenetic heterogeneity in the epiblast compared to more differentiated germ layers (Figure 5A), as well as a moderate correlation between promoter-wide DNA methylation-based and ATAC-based epiCHAOS scores from the same cells (Figure 5B).

**Figure 5.**
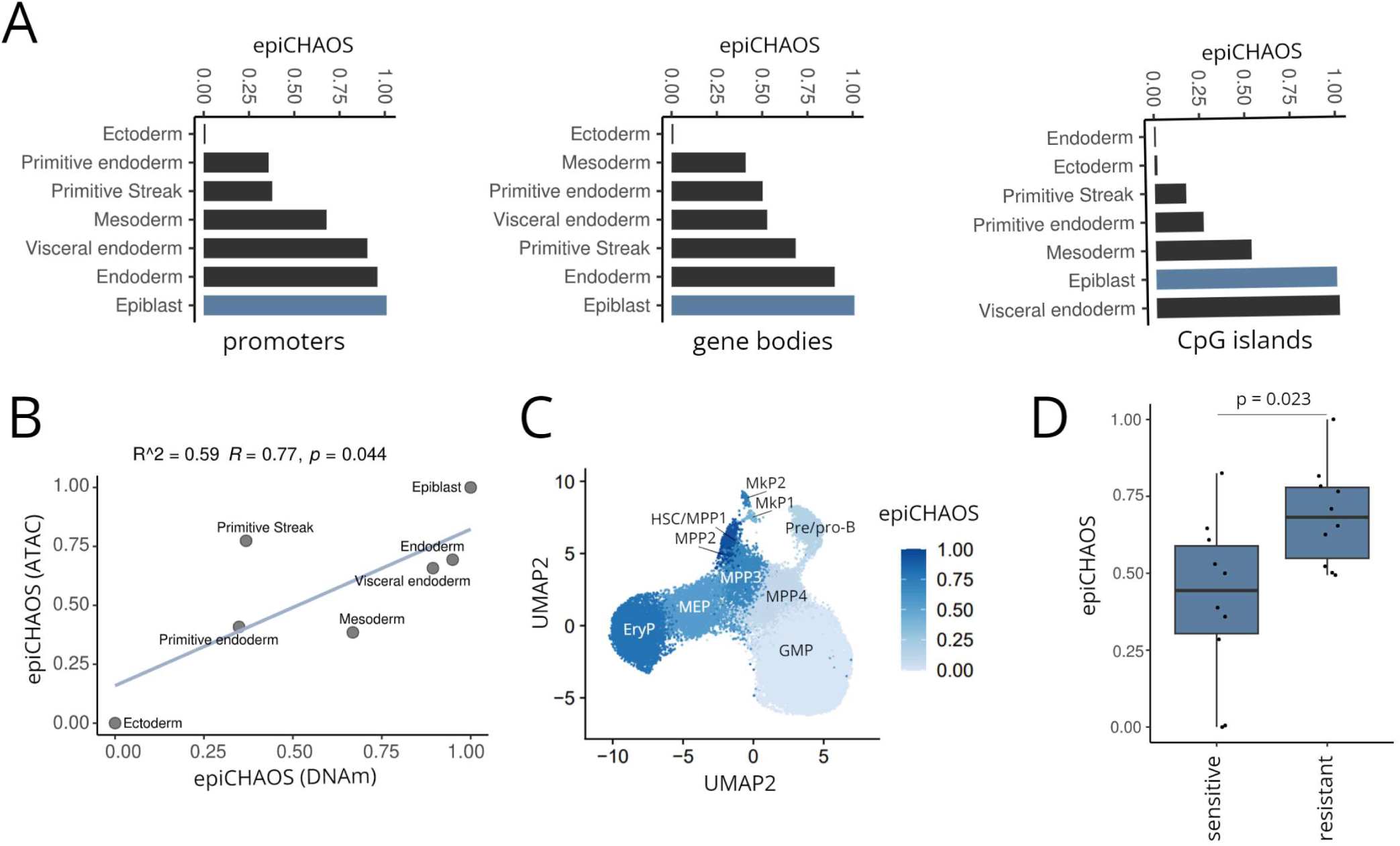
EpiCHAOS is applicable to single-cell epigenomics data from any modality. **A.** Barplots comparing epiCHAOS scores (epiCHAOS) across different lineages of mouse gastrulation using scNMT-seq DNA methylation data from Argelaguet et al.^42^ Methylation data summarized across promoters, gene-bodies or CpG islands were used for epiCHAOS computation. Epiblast is colored in blue. **B.** Scatter plot comparing promoter-wide epiCHAOS scores across different gastrulation lineages using single-cell DNA methylation [epiCHAOS (DNAm)] and ATAC-seq [epiCHAOS (ATAC)] data from the same cells as in A. Linear regression line is shown with Pearson correlation coefficient and p-value. **C.** UMAP embedding generated from scTAM-seq DNA methylation data from Scherer et al.^44^. Hematopoietic progenitor state and epiCHAOS scores are annotated. GMP: granulocyte monocyte progenitor, EryP: Erythroid progenitor, MEP: myeloid/erythroid progenitor, MPP: multipotent progenitor, HSC: hematopoietic stem cell, MkP: megakaryocyte progenitor. **D.** EpiCHAOS scores calculated using scCHIP-seq data for H3K27me3 from Grosselin et al.^42,45^. Cells were derived from a PDX breast cancer model, separated based on sensitivity or resistance to Capecitabine. Ten subsamples of 100 cells each were taken per condition. Boxplot shows comparison of epiCHAOS scores (epiCHAOS) between sensitive and resistant cells.

Next we utilized single-cell Targeted Analysis of the Methylome (scTAM-seq)^43^ data from mouse hematopoiesis^44^. In line with our previous observations at the level of chromatin accessibility (Figure 2A), we found that DNA methylation-based heterogeneity was increased in more primitive hematopoietic progenitor cells such as HSCs and early MPPs compared to more differentiated progenitors such as GMPs and pre-B cells (Figure 5C). Notably, we found a continuous decrease of epiCHAOS scores along the myeloid differentiation trajectory (HSC, MPP3, MPP4, GMP), while erythroid progenitors had elevated levels of heterogeneity in line with the scATAC-seq data shown in Figure 2A.

Finally, applying epiCHAOS to H3K27me3 single-cell chromatin immunoprecipitation sequencing (scChIP-seq) data from a previous study of breast cancer therapy resistance^45^, we detected higher epigenetic heterogeneity in therapy-resistant compared to sensitive cells, supporting the idea that intratumor epigenetic heterogeneity may serve as a driver of treatment resistance (Figure 5D).

We conclude that epiCHAOS is applicable to all single-cell epigenomics data types and not restricted to scATAC-seq data. This proves its utility across a wide range of epigenomic studies and enables the comparison of heterogeneity in multi-modal datasets.

## Discussion

Single-cell sequencing technologies offer unparalleled possibilities for studying epigenomic heterogeneity in development and disease. However, few studies have tackled the question of how cell-to-cell heterogeneity can be quantified at the level of cell populations, beyond the presence of distinct cell states. For instance, with current metrics one is unable to rank clusters or groups of cells by their heterogeneity levels. Here we introduce epiCHAOS: a quantitative metric for intercellular epigenetic heterogeneity computed at the single-cell level. We demonstrate how epiCHAOS can be used to provide unprecedented insights into the patterns of epigenomic variation across a wide range of applications including development and malignancy.

The sparsity of single-cell epigenomics data makes it particularly challenging to analyze and interpret in a continuous form, and binarization has become standard practice as a way to minimize the effect of sparsity and to reduce overall noise. We therefore specifically designed epiCHAOS for application to binary data. A limitation of this approach is that information about quantitative differences in accessibility between cells is lost, such that certain patterns of biological heterogeneity might be obscured. Nevertheless, we have demonstrated the functionality of our method by applying epiCHAOS to various scATAC-seq datasets to investigate both global and region-specific heterogeneity, where our results are consistent with existing biological paradigms. Our findings across a range of developmental contexts align with the accepted notion that less differentiated and more developmentally plastic cell types have higher heterogeneity compared to more differentiated and functionally specialized ones – a phenomenon that has never been demonstrated in single-cell epigenomics data^9,29^. We also find that epigenetic heterogeneity is increased with age in the hematopoietic system, reflecting the expected stochastic decay of epigenetic regulation^40^. Moreover, we provide evidence to support epigenetic heterogeneity as a hallmark of plasticity in cancer, paving the way for more detailed pan-cancer analyses in this domain.

Our findings regarding the preferential heterogeneity of specific genomic regions – in particular, the elevated cell-to-cell heterogeneity of polycomb binding sites – are also in agreement with previous studies of transcriptional noise and DNA methylation-based heterogeneity. For example, Kar *et al.* have previously shown that PRC2 targets display higher cell-to-cell variation in gene expression, with a low transcriptional burst frequency giving rise to oscillations between on and off states over time^41^. Kumar *et al*.^8^ also demonstrated that polycomb target genes are heterogeneously expressed within colonies of pluripotent stem cells. This heterogeneity may be due to the association of PRC2 targets with bivalent chromatin states – characterized by the presence of both active and repressive histone marks. Faure *et al.*^46^ also previously showed that genes with bivalent chromatin exhibit higher transcriptional noise. Similarly, Feinberg & Irizarry have demonstrated that DNA methylation stochasticity is increased at developmental genes – many of which are also PRC2 targets^47^. Here we demonstrate that these patterns of variability can be observed at the epigenetic level between single cells, and that they appear not only to be the most heterogeneous regions, but also the regions that are most preferentially heterogeneous in stem compared to more differentiated cells. It is notable that PRC2 targeted regions are also known to exhibit increased epigenetic variation across multiple cancers, compared to normal tissues^48^. This supports the idea that epigenetic heterogeneity at these regions might be a driving force in tumor evolution and raises interesting questions about the frequent disruption of PRC2 complex components in cancer^49^.

Understanding how epigenetic heterogeneity is shaped throughout tumor evolution including initiation, progression, remission and relapse, and deciphering its role in the progression to metastasis and therapy response could have important translational implications. EpiCHAOS could be used for example to investigate how epigenetic heterogeneity influences therapeutic resistance, or whether metastatic cells are epigenetically more heterogeneous than primary tumors. Moreover, epiCHAOS might also inspire strategies for guiding single-cell clustering and evaluating the similarity/variability across clusters in single-cell epigenomics data. Our method should therefore be especially valuable to cancer researchers, particularly those interested in plasticity and stemness, where the role of epigenetic heterogeneity remains underexplored. Beyond these questions, epiCHAOS should also yield novel biological insights outside of the cancer field, for instance in developmental biology, aging and immunity, and in other disease states in which plasticity programs are activated, such as wound healing and fibrosis.

## Conclusions

EpiCHAOS provides a quantitative metric of cell-to-cell epigenetic heterogeneity, complementing single-cell epigenomics studies of cancer and development. We have shown that epiCHAOS offers an excellent approximation of stemness and plasticity in various developmental contexts, as well as in cancer. Comparison of epiCHAOS scores at different genomic regions highlighted increased heterogeneity of polycomb targets and developmental genes. EpiCHAOS is applicable to a variety of single-cell epigenomics data types including measurements of chromatin accessibility, DNA methylation and histone modifications.

## Materials and Methods

### Calculating epiCHAOS scores

Heterogeneity scores were calculated for each given cluster or otherwise defined group of cells using a binarized matrix of scATAC or other single-cell epigenomics data. First pairwise distances were calculated between all cells within the cluster using a chance-centered version of the Jaccard index which controls for differences in the relative number of ones and zeros ^26^. Afterwards, the mean of all pairwise distances per cluster was computed. To remove any further effect of sparsity a linear regression model of the raw heterogeneity scores was fitted against the total number of detected accessible peaks, averaged across cells in the respective cluster, and the residuals of this model were taken as the adjusted scores. Finally, the scores were transformed to an interval of 0-1, and subtracted from 1 to convert the similarity metric to a distance metric.

ScATAC-seq data from the Hep-1 liver cancer cell line ^50^ was used to test whether epiCHAOS scores correlate with measures of technical noise. Cells were stratified into bins (20 bins of 100 cells each) based on various quality control metrics: FRIP scores, TSS enrichment scores and nucleosome ratio, which were calculated using ArchR ^51^.

DNA CNAs were inferred in the Hep-1 cell line using epiAneuFinder ^52^ with a windowSize=100,000 and minFrags=20,000. To investigate the influence of CNAs on epiCHAOS scores, the most prominent examples of large subclonal copy number gains (gain on chromosome 5) and deletions (deletion on chromosome 13) were selected by visual inspection, and cells were stratified based on the presence or absence of each alteration. EpiCHAOS scores were calculated across peaks in the affected chromosome and compared between cells with diploid or alternative states. To correct for CNAs when applying epiCHAOS to cancer datasets, a per-chromosome count-corrected epiCHAOS score was derived, where epiCHAOS scores were calculated per chromosome, implementing a linear regression-based adjustment for the total coverage on that chromosome, and then the average of per-chromosome scores was taken as the global epiCHAOS scores.

Unless otherwise specified, epiCHAOS was calculated using the entire peaks-by-cells matrix. To allow a more robust comparison between groups, epiCHAOS scores were calculated on five random subsamples of 100 cells from each group/cluster, except in groups/clusters which contained fewer than 100 cells. Since the scNMT-seq data contained fewer than 100 cells in most groups, epiCHAOS scores were calculated only once for each cell type. ENCODE Transcription Factor Binding Site (TFBS) regions from the LOLA core database^53^ were used for comparisons of heterogeneity at different genomic regions, for which scATAC peaks matrices were subsetted to obtain peaks overlapping with each genomic region. Similarly for comparisons across gene sets, data were subsetted for peaks overlapping with promoters of each gene set using the GO:BP gene sets from MsigDB^54^.

### Generating synthetic datasets with controlled heterogeneity

To test the performance of epiCHAOS, synthetic datasets were generated *in silico* in a way that simulates the structure of binarized scATAC-seq peak matrices. First a series of 100 synthetic datasets with controlled heterogeneity was created, in which each dataset has an equal total count. To do this a random binary matrix was created, which would represent the first dataset in the series. In each subsequent dataset, homogeneity was incrementally introduced by removing a set number of 1’s from selected n rows, and adding them to a different selected n rows, in such a way that a constant number of 1’s is maintained, while heterogeneity decreases.

To test situations where the genome-wide chromatin accessibility is increasing or decreasing, binarized data from an example scATAC-seq dataset were perturbed to create datasets of increasing heterogeneity with either addition or removal of 1’s. Specifically, 10, 20, 30, 40 and 50% of 1’s were selected at random and replaced by 0’s, and corresponding numbers of 0’s were selected at random and replaced by 1’s.

To test that epiCHAOS is not influenced by differences in sparsity, a series of 100 random binary datasets was generated with each dataset having equal dimensions and incrementally increasing total number of 1’s. Their epiCHAOS scores were then computed and tested for a correlation with their total count.

As an additional validation approach semi-synthetic scATAC-seq datasets were created by mixing data from distinct cell types. Using scATAC-seq data from human bone marrow ^28^ five cell types were selected; HSCs, Monocytes, CD8-CM T cells, B-cells and plasmacytoid DCs. The top 500 differentially accessible peaks for each cell type were identified and used to create *in silico* mixtures of two to five cell types in all possible combinations.

### Simulating scATAC-seq data with varying sequencing depth

The scReadSim package was used to simulate scATAC-seq data of varying sequencing depths ^27^. A subset of scATAC-seq data from HSCs from the Granja et al. dataset was used as input^28^. Data were reduced to 10,000 randomly selected peaks for ease of processing. Simulated scATAC-seq matrices comprising each 500 cells were generated with sequencing depth ranging from 50,000 to 100,000 counts, in increments of 10,000. EpiCHAOS scores were calculated across matrices on five subsamples of 100 cells from each condition.

### scATAC-seq data processing and analysis

Publicly available scATAC-seq datasets for human hematopoiesis ^28^, mouse gastrulation ^30^, drosophila embryogenesis ^55^, breast cancer ^13^, liver cancer ^34^, ependymoma ^36^, HSC aging ^39^ and liver cancer cell lines ^50^ were downloaded from the respective publications. For analyses in developmental datasets and in ependymoma processed counts matrices were used as provided by the authors, where cell types were previously annotated. For breast and liver cancer datasets fragments files were downloaded, processed and analyzed using ArchR^51^. Cells with TSS enrichment scores less than 4 or number of fragments higher than 1,000 were removed, and doublets were filtered out using default parameters. Iterative LSI was performed followed by clustering using the Seurat method. Gene score matrices were generated using ArchR and used for subsetting cancer datasets for epithelial cells based on epithelial cellular adhesion molecule (EPCAM) scores. After reclustering epithelial cells, peak calling was performed using macs2^56^. To assign gene set/pathway scores to each cluster, gene set annotations were obtained from MSigDB using the msigdbr R package^54^. Gene scores were first averaged across all cells within each cluster, and then the mean score of all genes within a given gene set was computed to assign gene set scores per cluster.

### Differential heterogeneity analysis

Differential heterogeneity analyses were performed for each region using a permutation approach, whereby the difference in epiCHAOS scores between two selected cell types were compared with that between pairs of 1,000 randomly computed groups sampled from the same pool of cells. P-values were calculated as the quantile of the distribution of sampled permutations for which the difference in heterogeneity scores was greater than that between the two test groups.

### DNA methylation variation

Data from Adelman et al. 2019 ^57^ was used to calculate DNA methylation variability by computing variance per CpG site in HSC-enriched lineage-negative (Lin− CD34+ CD38−) samples across the eight male donors. To address the issue of data sparsity, a maximum quantile threshold of 0.005 was established for missing values per site. Any sites that surpassed this threshold were removed. For each ENCODE TFBS, the average of variances was calculated for all CpG sites overlapping with the respective regions.

### scRNA-seq analyses

Publicly available scRNAseq datasets for human hematopoiesis^28^, mouse gastrulation ^30^ and drosophila embryogenesis^55^ were downloaded from the respective publications and analyzed using Seurat. Cell-to-cell transcriptional heterogeneity was calculated by computing pairwise euclidean distances according to the methods of Hinohara *et al.* ^19^. Developmental potential was calculated per cell using CytoTRACE ^33^ and assigned as a mean per celltype for downstream analyses. Transcriptional noise per gene was estimated using the coefficient of variation as previously described ^58^. A list of PRC2-target genes used for comparison of transcriptional noise was obtained from Ben-Porath *et al.* ^59^

### scCHIP-seq analysis

scCHIP-seq counts matrices representing 50kb non-overlapping bins of H3K27me3 from human breast cancer patient-derived xenograft cells that were sensitive or resistant to Capecitibine (HBCx-95 and HBCx-95-CapaR) were downloaded from GSE117309 and processed as described by Grosselin et al. ^45^. Cells having a total number of unique reads in the upper percentile were removed as outliers, and genomic regions not represented in at least 1% of all cells were filtered out. Data corresponding to non-standard chromosomes and the Y chromosome were excluded. Cells with a total number of unique reads less than 1,600 were removed. Counts matrices were binarized and cells from each condition were subsampled to select ten groups of 100 cells each for epiCHAOS calculation.

### scNMT-seq and scTAM-seq analysis

scNMT-seq DNA methylation and ATAC data from mouse gastrulation ^42^, summarized across promoters, gene bodies and CpG islands, were accessed using the SingleCellMultiModal R package. scTAM-seq data from mouse hematopoiesis were obtained from Scherer et al.^44^, downloaded from https://figshare.com/ndownloader/files/42479346, and analyzed using Seurat.

## Declarations

### Ethics approval and consent to participate

Not applicable

### Consent for publication

Not applicable

### Availability of data and materials

All datasets analyzed in this article are publicly available and can be accessed through their associated publications as mentioned in the “Methods” section. EpiCHAOS is available as an R package at CompEpigen/epiCHAOS (github.com). Analysis scripts used throughout the manuscript are deposited on GitHub at CompEpigen/epiCHAOS-manuscript (github.com).

### Authors’ contributions

KK developed the method, devised and performed the majority of analyses. CP proposed the project idea and supervised the work. PL co-supervised the work and proposed strategies for assessing differential heterogeneity and accounting for copy number alterations. MS contributed to the overall design and planning of the manuscript, proposed the comparison with DNA methylation-based heterogeneity and the application of the method to single-cell DNA methylation data. MB performed the calculation of DNA methylation variation. KK wrote the manuscript and CP, MS and PL all discussed and contributed to its final version.

## Acknowledgements

We would like to thank Karsten Rippe, Isabelle Seufert, Sebastian Waszak, Moritz Mall, Manu Saraswat, and members of the division of Cancer Epigenomics at the DKFZ for insightful discussions about the method. We are also thankful to Thomas Hielscher for statistical recommendations, and Dieter Weichenhan for comments on the manuscript.

