## Supplementary Figures for "EpiCHAOS: a metric to quantify epigenomic heterogeneity in single-cell data"

### Additional file 1: Supplementary Figures

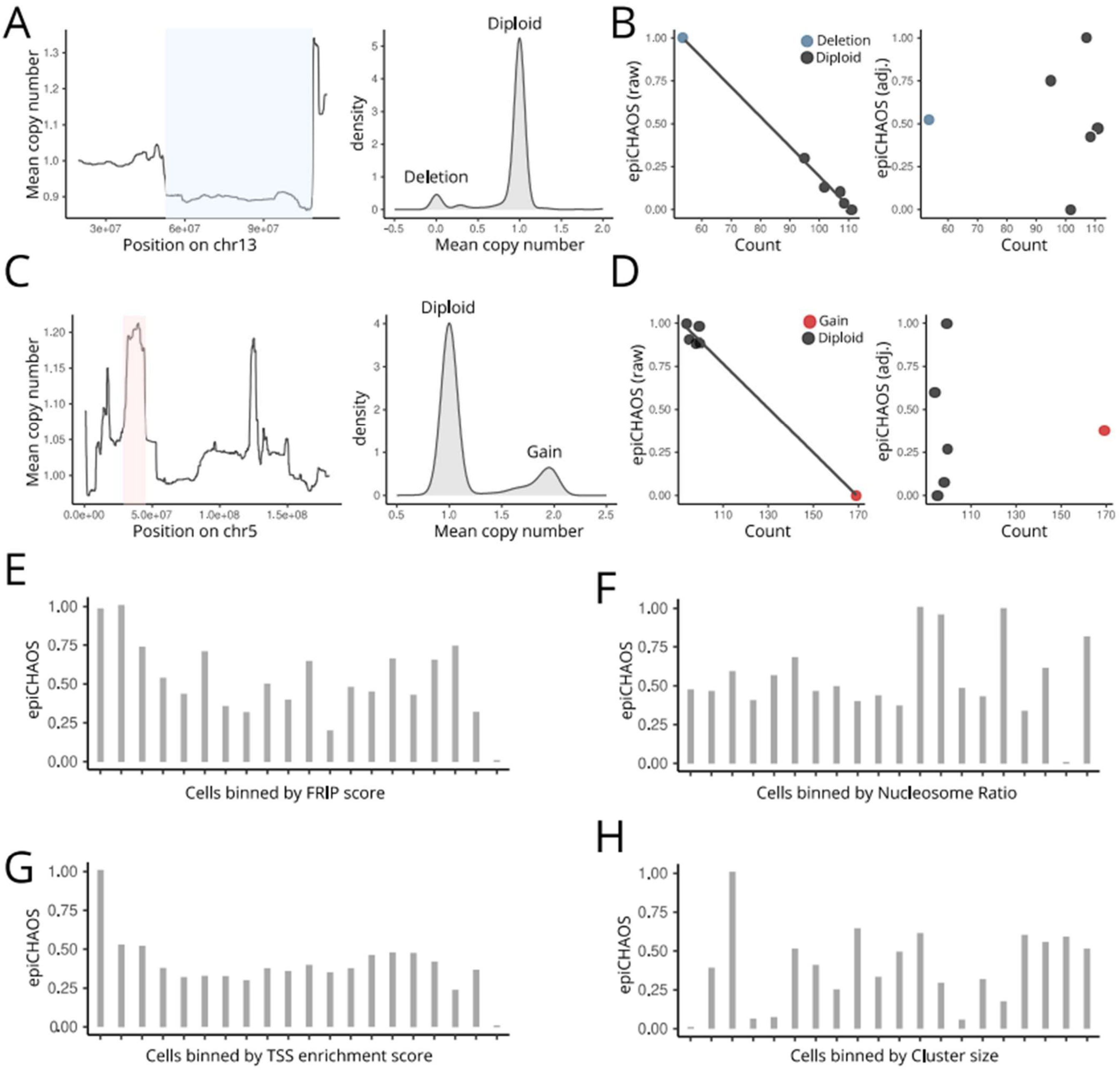


**Figure S1. EpiCHAOS scores are not influenced by differences in copy number or technical confounders**

**A.** Selection of cells with subclonal copy number loss on chromosome 13 in the Hep-1 cell line. Line plot (left) shows the mean copy number among single cells on chromosome 13. Copy number was called using epiAneuFinder, where 1 = diploid, 0 = loss and 2 = gain. X-axis displays chromosome location. The highlighted region was selected as a region of subclonal deletion for which to test the effects of copy number alterations on epiCHAOS scores. Density plot (right) shows the distribution of cells with and without deletion in the selected region. **B.** After reducing the peaks-by-cells matrix for peaks within the deleted region (highlighted in (A)), one group of 100 deletion cells and five groups of 100 diploid cells were sampled for epiCHAOS calculation. Scatterplots show the relationship between epiCHAOS scores (epiCHAOS) and counts (average counts per cell in the group in the subsetted peaks matrix) before (left) and after (right) adjustment for total counts. EpiCHAOS scores were adjusted for counts by fitting a linear regression model of epiCHAOS scores against counts (average counts per cell in the group) and taking the residuals of the model as an adjusted score. Each point represents a group of either diploid or deleted cells on which epiCHAOS scores are computed. **C.** Selection of cells with subclonal copy number gain on chromosome 5 in the Hep-1 cell line. **C-D.** Plots show the same as in (A-B) in the situation of a copy number gain (increased counts). **E.** Bar plot of epiCHAOS scores (epiCHAOS) computed on 20 bins of 100 Hep-1 cells ordered by increasing FRIP scores. **F.** Bar plot of epiCHAOS scores (epiCHAOS) computed on 20 bins of 100 Hep-1 cells ordered by increasing TSS enrichment score. **G.** Bar plot of epiCHAOS scores (epiCHAOS) computed on 20 bins of 100 Hep-1 cells ordered by increasing nucleosome ratio. **H.** Bar plot of epiCHAOS scores (epiCHAOS) computed on 20 groups of randomly selected Hep-1 cells, where the number of cells in each group increases in increments of 25 cells, from 25 to 500 cells.

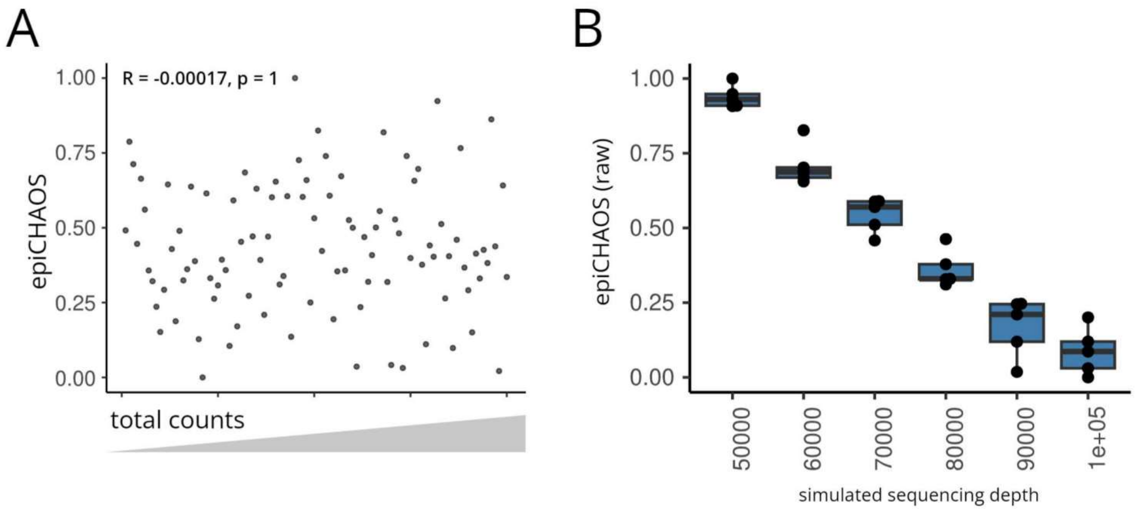


**Figure S2. Assessing the effect of sparsity and sequencing depth on epiCHAOS scores**

**A.** Scatterplot illustrating the absence of correlation between epiCHAOS scores (epiCHAOS) and total coverage across a series of 100 randomly generated datasets ordered by increasing total counts. Pearson correlation coefficient and p-value is shown. **B.** Boxplot comparing raw epiCHAOS scores (before adjustment for sparsity) across six simulated single-cell ATAC-seq datasets. Data were simulated using scReadSim with sequencing depth varying from 50,000 to 100,000 counts. ScATAC-seq data from hematopoietic stem cells subset from the Granja *et al.* 2019 dataset were used as the baseline counts matrix.

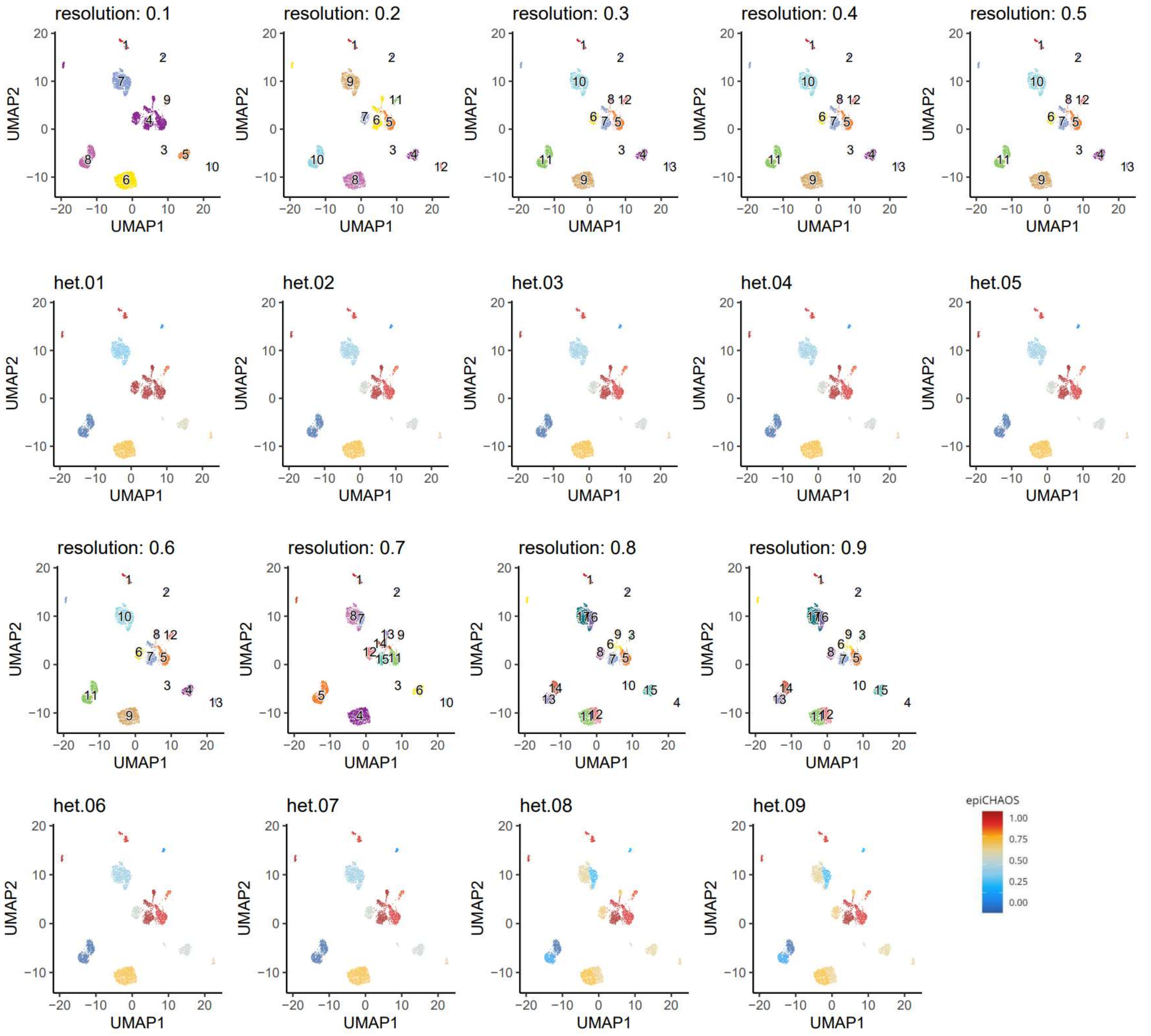


**Figure S3. EpiCHAOS scores are minimally influenced by clustering parameters**

UMAP embeddings of breast cancer epithelial cells using scATAC-seq data from Kumegawa et al. Top row UMAPs show the clustering result from ArchR using different clustering resolutions from 0.1 to 0.9, as labelled. Bottom row UMAPs show the epiCHAOS scores (epiCHAOS) calculated for each cluster using the respective resolution.

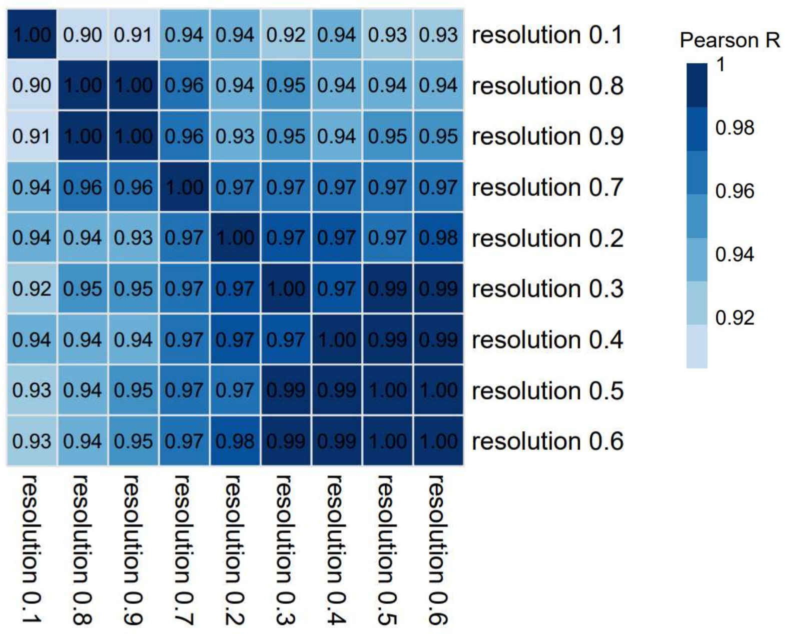


**Figure S4. Correlation of epiCHAOS scores across single cells at different clustering resolutions**

Heatmap displays the Pearson correlation coefficients from per-single-cell correlation of epiCHAOS scores across different clustering resolutions as depicted in Figure S3.

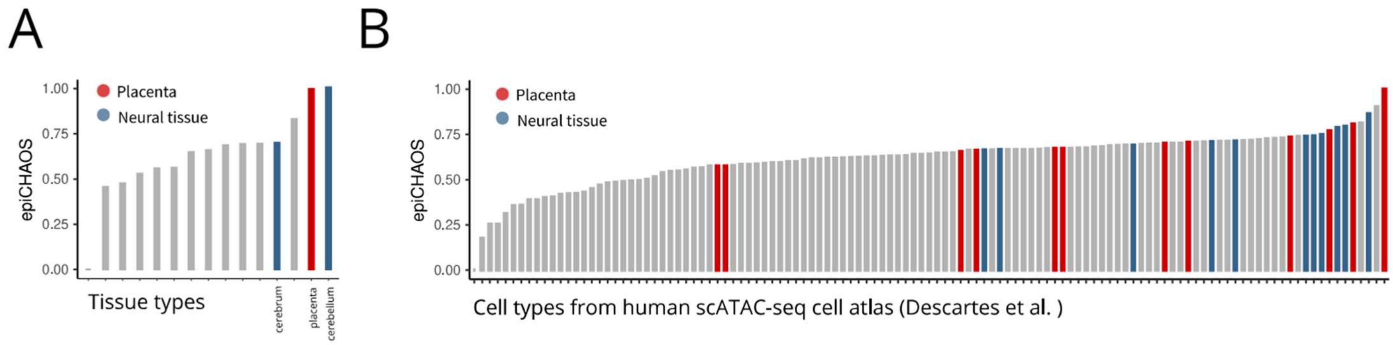


**Figure S5. Epigenetic heterogeneity is increased in developmental and neuronal tissues**

**A-B.** Barplots of epiCHAOS scores (epiCHAOS) in different tissues (A) and cell types (B) from the human scATAC-seq cell atlas (Descartes et al.). Neural and placental tissues are highlighted.

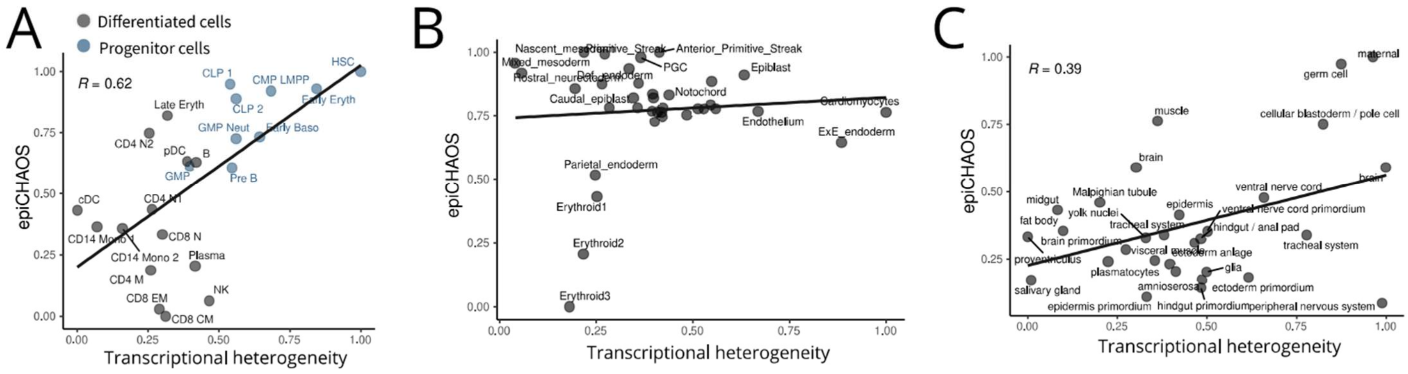


**Figure S6. Comparison of epigenetic and transcriptional heterogeneity in developmental settings**

**A-C.** Scatterplots show the correlation of epiCHAOS scores (epiCHAOS) with transcriptional heterogeneity scores in human hematopoiesis (A), mouse gastrulation (B) and drosophila embryogenesis (C). Transcriptional heterogeneity is quantified using the associated scRNA-seq data (not data from the same cells) by taking the mean of pairwise euclidean distances between all cells within a group/cluster.

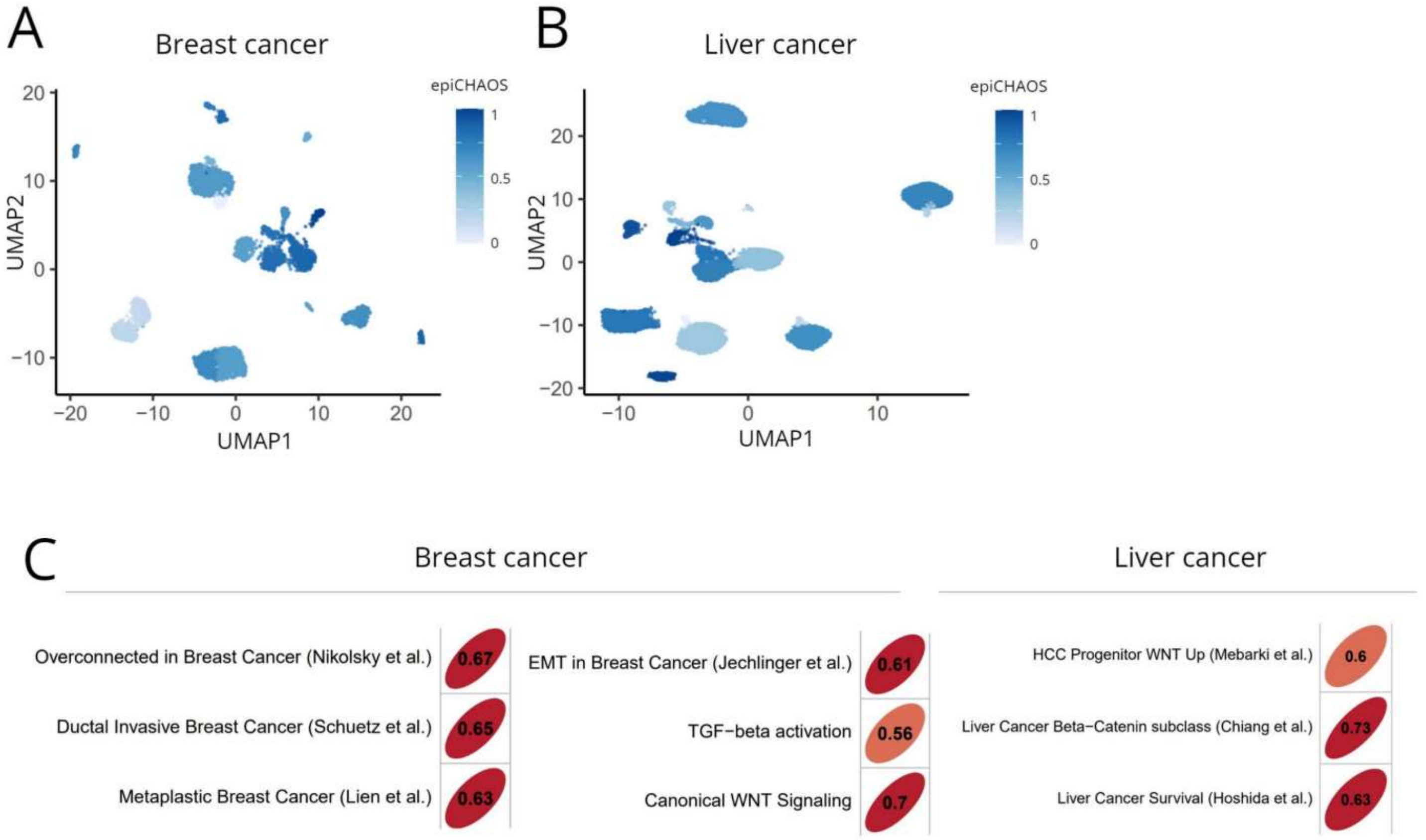


**Figure S7. Application of epiCHAOS to scATAC-seq data from breast and liver cancer**

**A-B.** UMAP representations of scATAC-seq clusters from (A) breast and (B) liver cancer samples after subsetting epithelial cells. Clusters are colored by epiCHAOS scores (epiCHAOS). **C.** Correlation plots of epiCHAOS scores with selected gene sets from MSigDB. Pearson correlation coefficients are shown.

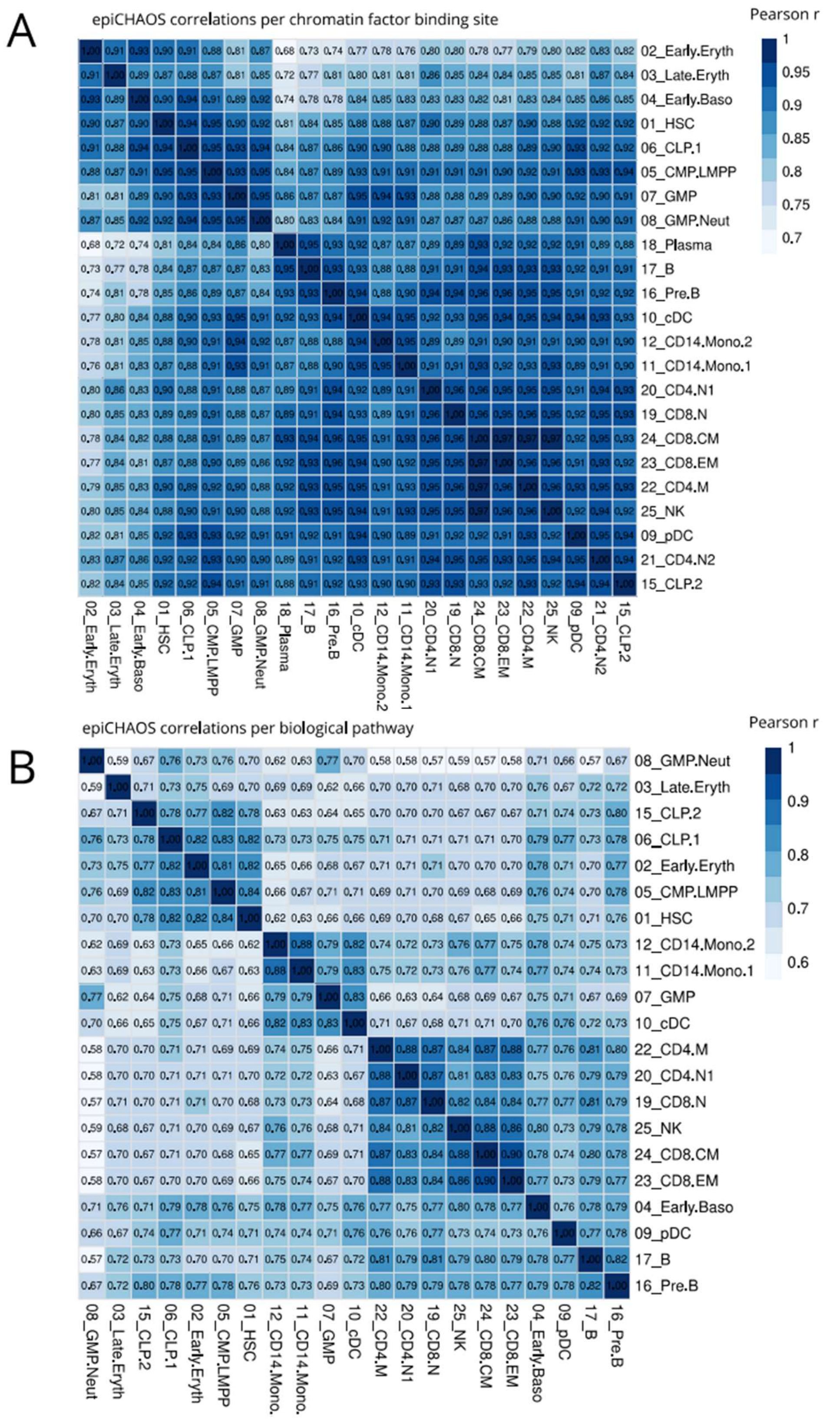


**Figure S8. Patterns of heterogeneity at different genomic regions are correlated across hematopoietic cell types**

**A-B.** Correlation heatmaps depicting the similarity in per-region and per-gene set epiCHAOS scores between cell types from human bone marrow (Granja et al.). Correlations were computed across all (A) chromatin factor binding sites from the ENCODE TFBS database and (B) gene ontology biological processes. Pearson's correlation coefficients are shown.

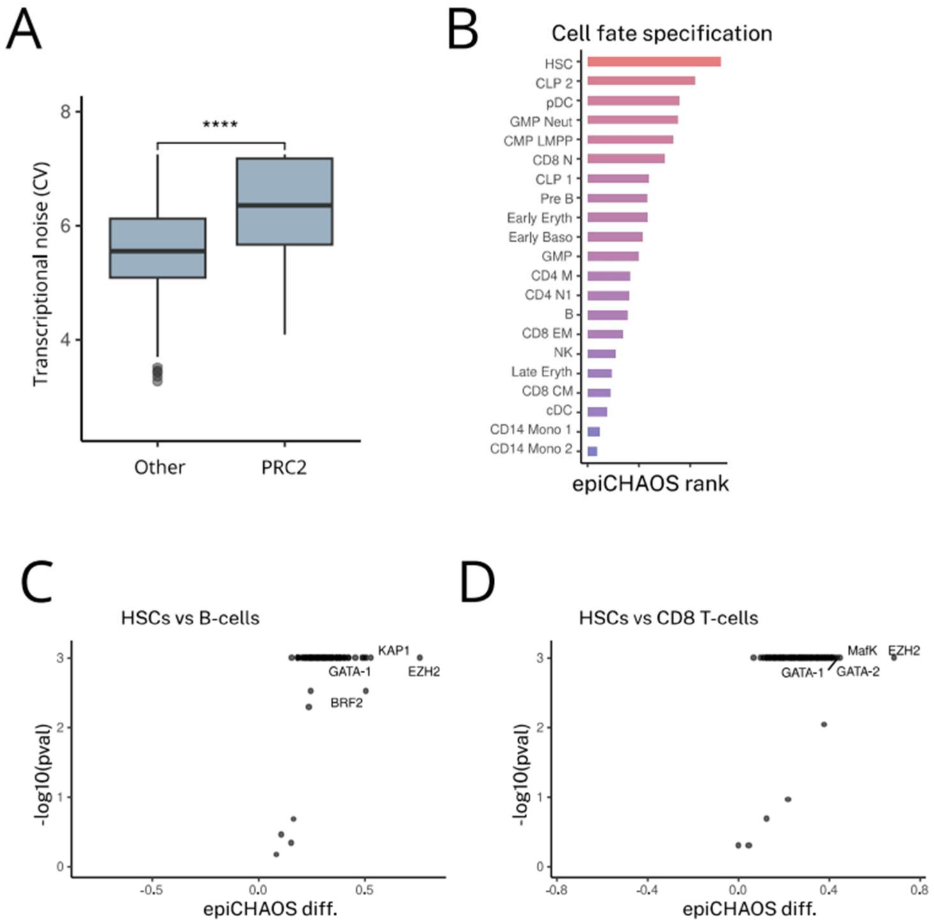


**Figure S9. Epigenetic heterogeneity is increased at PRC2 targets and developmental genes**

**A.** Boxplot comparing transcriptional noise, measured using the coefficient of variation, between PRC2 target genes and other genes in hematopoietic stem cells using scRNA-seq data from Granja et al. **B.** Barplot comparing epiCHAOS ranks for the gene ontology biological process "cell fate specification" across different hematopoietic cell types. The higher the rank indicates that the selected gene set has higher epiCHAOS scores compared to other gene sets in that cell type. Ranks were -log(10) transformed for display. **C-D.** Volcano plots illustrate differential heterogeneity between hematopoietic stem cells and B cells (C) and CD4 Memory T-cells (D). Differential heterogeneity was tested for each ENCODE TFBS (binding sites from K562 cells). For each TFBS, the-log10(p-value) obtained by permutation test is displayed on the y-axis, and the difference in epiCHAOS scores between the two cell types is displayed on the x-axis.
