## Supplementary Notes for "EpiCHAOS: a metric to quantify epigenomic heterogeneity in single-cell data"

### Additional file 2:

#### Supplementary Note 1

To make sure that "noisier" data does not result in a higher heterogeneity score, we assessed the relationship between epiCHAOS scores and a range of commonly used metrics of technical noise in scATAC-seq data; the fragments of reads in promoters (FRIP) score, transcription start site (TSS) enrichment score and nucleosome ratio. EpiCHAOS scores did not correlate with any of the above metrics of cell quality except in cases of extremely low TSS enrichment scores which would normally be filtered out as part of standard quality control (Additional file 1: Figure S1E-G). In any case, since the heterogeneity score provides a cluster-level, rather than cell-level statistic, it should not be affected by technical noise as long as we are comparing cells profiled within the same experiment. EpiCHAOS was also not influenced by the number of cells per cluster (Additional file 1: Figure S1H).

#### Supplementary Note 2

To investigate how epiCHAOS behaves under different clustering parameters, we selected as an example the breast cancer dataset from Kumegawa et al. which we used in Figure 3. We performed clustering in ArchR using a range of clustering resolutions from 0.1 to 0.9. We then computed epiCHAOS scores at each different clustering resolution and compared the results. While we do find that splitting a large cluster into several smaller clusters can reveal which parts of that larger cluster are more heterogeneous than others, the overall results change minimally. We find that a similar pattern of epiCHAOS scores emerges in different cell groups regardless of clustering resolution (Figure S3), with a high correlation of per-single cell epiCHAOS scores measured at different resolutions (Figure S4). This suggests that epiCHAOS scores are minimally influenced by the choice of clustering parameters.
